# Costs and benefits of giant sperm and sperm storage organs in *Drosophila melanogaster*

**DOI:** 10.1101/652248

**Authors:** Susanne Zajitschek, Felix Zajitschek, Sarah Josway, Reem Al Shabeeb, Halli Weiner, Mollie K. Manier

**Affiliations:** George Washington University, Department of Biological Sciences, Science and Engineering Hall, 800 22nd St. NW, Suite 6000, Washington, DC 20052, United States; School of Biological, Earth and Environmental Sciences, University of New South Wales, Kensington Campus, Sydney, NSW 2052, Australia

**Author notes:** Joint first authors (Susanne Zajitschek, Felix Zajitschek). Correspondence: Mollie Manier.

**Keywords:** fecundity, Fisherian runaway selection, good genes, intralocus sexual conflict, longevity, mating success, pre-copulatory sexual selection, post-copulatory sexual selection, sperm, sperm storage organ

## Abstract

In the *Drosophila lineage*, both sperm and the primary female sperm storage organ, the seminal receptacle (SR), may reach extraordinary lengths. In *D. melanogaster*, long SRs bias fertilization toward long sperm during the displacement stage of sperm competition. This sperm-SR interaction, together with a genetic correlation between the traits, suggests that the coevolution of exaggerated sperm and SR lengths may be driven by Fisherian runaway selection. To further understand the costs and benefits of long sperm and SR genotypes in both sexes, we measured male and female fitness in inbred lines of *D. melanogaster* derived from four populations previously selected for long sperm, short sperm, long SRs, or short SRs. We specifically asked: do long SRs impose costs or benefits on the females that bear them? Do genotypes that generate long sperm in males impose a fitness cost on females sharing those genotypes? Is long sperm an honest indicator of male viability and associated with increased fitness? And finally, are the benefits of long sperm restricted to competitive fertilization success, or do long-sperm males also have increased mating success and fecundity in single matings? We found that both sexes have increased longevity in long sperm and long SR genotypes, with fewer reproduction-related benefits and evidence for trade-offs in males, compared to females. Our results suggest that sperm length and SR length are both indicators of increased viability.

## INTRODUCTION

Post-copulatory sexual selection can drive the rapid evolution of male ejaculate traits across diverse taxa, mediated by sperm competition (Parker, 1970) on the one hand and cryptic female choice (Eberhard, 1996; Firman et al., 2017) on the other. These two processes occur after mating in an analogous fashion to male-male competition and female choice, which comprise pre-copulatory selection that acts before mating. Male traits under pre-copulatory sexual selection often take the form of elaborate visual, audible, tactile, and/or chemical displays, and female preferences for them are based on sensory perception that leads to behavioral decisions (Candolin, 2003; Jennions and Petrie, 1997). Female preference under post-copulatory sexual selection is called cryptic female choice and occurs when female-mediated behavioral, morphological, or physiological processes bias paternity in favor of certain males (Pitnick and Brown, 2000), based on pre-copulatory (Pilastro et al., 2004; Sbilordo and Martin, 2014) or post-copulatory male traits (Wojcieszek and Simmons, 2012). Whether acting before or after copulation, female preference evolution follows similar expectations predicted under runaway selection (Fisher, 1958; Kirkpatrick, 1982), good genes (Iwasa and Pomiankowski, 1991; Zahavi, 1975), or sexy son (Pomiankowski et al., 1991)/sexy sperm (Keller and Reeve, 1995) hypotheses.

In the *Drosophila* lineage, sperm reach extraordinary lengths (Scott Pitnick et al., 1995), which is presumably driven by post-copulatory sexual selection, and mediated by the length of the female’s primary sperm storage organ, the seminal receptacle (SR), which can be even longer (Pitnick et al., 1999). Long sperm have a competitive fertilization advantage against shorter sperm, but primarily within long SRs (Miller and Pitnick, 2002). Specifically, sperm competition between experimentally evolved long sperm and short sperm populations of *D. melanogaster* revealed a competitive advantage of long sperm but only in experimentally evolved females with long SRs. This long sperm advantage occurs through as yet undescribed fluid dynamic processes during the displacement stage of sperm competition (Manier et al., 2010; 2013) that allows longer sperm to retain a positional advantage closer to the site of fertilization (Pattarini et al., 2006). Thus, variation in SR length is a proxy for the strength of cryptic female choice for sperm length, with longer SRs being more selective, or “choosier”, based on the size of the post-copulatory male ornament, sperm length.

If *Drosophila* sperm length is a male ornament, a number of patterns could be expected; (1) If this exaggerated trait has evolved under runaway selection, the male ornament and female preference should coevolve together and be genetically correlated. (2), if long sperm carry indirect benefits consistent with a good genes model of ornament evolution, they should also be costly and condition-dependent (Zahavi, 1977), and possibly trade off with other male traits (reviewed in Manica et al., 2016). Finally (3), we could expect long sperm to display strong positive allometry (Bonduriansky, 2007; Kodric-Brown et al., 2006; Voje, 2016), particularly if sperm length could be considered a “weapon” rather than a “display” (Eberhard et al., 2018). In support of these predictions, sperm length and SR length are coevolving both among species (Pitnick et al., 1999) and among populations within *D. mojavensis* (Pitnick et al., 2003), and there is a significant genetic correlation between the two traits (Lüpold et al., 2016). Long sperm are also costly in terms of time required to reach reproductive maturity (Miller and Pitnick, 2002; Pitnick et al., 2003; Pitnick et al., 1995), and sperm length trades off with sperm number across species (Pitnick, 1996). Moreover, condition-dependence of sperm length increases in species with longer sperm (Lüpold et al., 2016), and as expected for certain male ornaments, sperm length has the strongest positive allometry with body size ever measured for a sexually selected trait (Lüpold et al., 2016).

Here, we use *Drosophila* sperm length and SR length as a model to investigate trait-preference coevolution. Specifically, we ask whether long sperm and SRs carry additional fitness costs or benefits outside of post-copulatory sexual selection. Fitness consequences of exaggerated trait genotypes manifested in either sex could influence the dynamic of sperm-SR coevolution, either by reinforcing selection in the same direction on both sexes or imposing an antagonistic relationship between selection on males and females. This system has a unique advantage in that the female “preference” (SR length) is an easily and consistently quantifiable morphological trait, rather than a behavioral or cognitive process that may be more difficult to measure and is potentially affected by social learning. In order to better understand the nature of male-female coevolution, we need to elucidate the costs and benefits of male ornament and female choice genotypes for both the sex in which they are expressed and the sex in which they are not. Long sperm have a number of costs, as outlined above, and it is yet unclear if their post-copulatory competitive advantage also transfers to increased attractiveness, mating success, or fertility. Sperm size could be correlated with these traits due to genetic linkage with viability alleles (Gilbert and Uetz, 2016; Head et al., 2005; Svobodová et al., 2018), have significant trade-offs (Ball and Parker, 1996; Dines et al., 2015; Foo et al., 2018), or be uncorrelated (Travers et al., 2016), depending on a range of ecological factors (Evans and Garcia-Gonzalez, 2016; Lüpold et al., 2014; Parker et al., 2013; Simmons et al., 2017).

Similarly, long SRs are associated with longer development times but increased fertility (Miller and Pitnick, 2003, 2002). However, it remains unclear whether genotypes that produce long sperm or long SRs confer similar costs and benefits when present in the opposite sex. The ability of female preference to drive the evolution of exaggerated male traits must be robust to moderate the costs of female choice (Chandler et al., 2013; Mead and Arnold, 2004), but the strength and direction of male-female coevolution may be affected by fitness consequences incurred by trait genotypes in the sex not expressing the trait; (Chenoweth et al., 2008; Chippindale et al., 2001; Cox and Calsbeek, 2009, Pischedda and Chippindale, 2006).

We investigated benefits and trade-offs associated with sperm length or SR length genotypes for both males and females in populations previously selected for sperm or SR length (Miller and Pitnick, 2002). Specifically, we measured male mating success, male and female fertility, and male and female longevity in isofemale lines with long sperm, short sperm, long SRs, or short SRs. We found sex-specific trade-offs for both long sperm and long SRs with mating success and fecundity, respectively. However, long sperm genotypes did confer longevity benefits for both males and females, and for males from long SR lines. We did not detect any direct fitness benefits for long SR females, but males from long SR lines had higher fecundity. Taken together, these results suggest that male-female coevolution of sperm length and SR length in this system is facilitated by both increased viability and indirect benefits of long sperm and SRs in both sexes. That is, long SR females and long sperm males lived longer (viability benefits), and by selecting for long sperm, long SRs in females may provide indirect benefits through increased longevity in both sons and daughters.

## METHODS

### Experimental populations

To determine fitness effects of sperm length or SR length, we quantified mating success, fecundity, and longevity in four *D. melanogaster* populations that had been previously selected for long sperm, short sperm, long SRs, or short SRs (initially reported in Miller and Pitnick, 2002). Initially, a panel of inbred isolines was created by mating full siblings for ten generations. Within each of the four selection regimes, the four most extreme isolines were identified and used for this experiment. For each isoline, a minimum of five female SRs, and, on average 5 sperm cells (range: 2-11 sperm) from each of at least four males (range 4-8, average: 5.56) were measured. To measure sperm, seminal vesicles from mature virgin males (5 days post-eclosion) were dissected into a large droplet of 1X phosphate buffered saline (PBS) on a glass slide, ruptured, and dragged several times to release the live sperm. The droplet was dried down at 55 C, and sperm were fixed in 3:1 methanol:acetic acid, mounted in glycerol, and the slide sealed with nail polish. Sperm were visualized on a Nikon Ni-U upright light microscope at 200X magnification under darkfield. Images were captured with an Andor Zyla 4.2 camera and measured using the segmented line tool in ImageJ (https://imagej.nih.gov/ij/).

SRs were measured from mature virgin females (5-7d post-eclosion) that were stored frozen (−20°C) until dissection. Female reproductive tracts were dissected into 1X PBS, the SR gently unraveled with a fine insect pin, and the sample mounted under a coverslip, such that the SR was two-dimensional but not over-compressed. SRs were visualized at 100X magnification under phase contrast, and images were captured and measured as outlined above.

From across three experimental blocks, a total of *N* = 1151 males and *N* = 1298 females were included in the final analyses. All stocks were maintained at ambient room temperature and light regime in polyethylene fly vials with cornmeal agar yeast molasses medium supplemented with live yeast. Experimental flies were reared by pairing 2-3 day old female and male flies from the same isoline for 48 hours, and emerging offspring were collected as virgins. All reproductive and behavioral assays were performed at the same time of day to reduce circadian rhythm effects. All individuals were collected as virgins under light CO_2_ anesthesia, maintained in same-sex vials with densities of 10 females or 20 males, and were 2-5 days old when first mated.

### Mating success

We observed male mating behavior to assess attractiveness (latency to mate), copulation duration, and the proportion of successful matings for males from each of the four selection regimes: long sperm, short sperm, long SR, and short SR. A subset of ten randomly selected males from each of three replicate vials from each of the 16 isolines (*N* = 460) were tested for five consecutive hours (or until successful mating) each week over a period of six weeks. Individual males were transferred without anesthesia to vials with a single 5-day-old wild type (LHm) virgin female into a mating arena consisting of a polyethylene vial with a foam plug in the bottom to enhance visibility, and a cotton plug pushed halfway down the vial to stimulate male-female interactions. For each successful mating, latency to mate and mating duration were recorded, after which males were returned to their original vial. Males were transferred to new food vials three times a week, and dead males were removed without replacement, with date of death recorded for longevity analyses.

### Fecundity

To measure female fecundity, experimental virgin females 2-3 days post-eclosion were paired individually with a wild type LHm male (5 to 7 days old) for 48 hours, after which the male was removed (*N* = 160). Each week, we subsampled progeny produced within a 24 hr period for each female over the course of her life (see Longevity, below). Specifically, we allowed the eggs that had been laid within the specified weekly 24 hr period to develop, and counted the number of eclosed and uneclosed pupae, four days after the flies in a given vial had started hatching. All weekly counts from each female were summed to approximate lifetime reproductive output.

For male fecundity, we counted progeny produced by up to two randomly selected successfully mated males from each replicate vial (N max/week = 96) for each week of the mating success assays (see above). Specifically, LHm females were separated from the males directly after mating, and transferred to a new food vial, where they were allowed to deposit eggs for 48 hours before being discarded. Adult offspring were counted as a proxy for male fecundity. In contrast to females, male offspring data were not measured on the same individuals over time, as the individual identity of males within a given vial was unknown.

### Longevity

Males were kept in cohorts of initially 20 same-sex flies per vial (three replicate vials per isoline, populated one day post-eclosion, *N* = 48 vials). We checked for survival every two days, when flies were transferred to a new food vial. We tested how selection regime affected survival using Cox proportional hazard models (function *coxph*; Therneau, 2015), separately for each sex and each selected trait. Females were maintained individually to assess female reproductive success (10 replicates each, conducted in blocks 1 & 2).

### Statistical Analyses

To analyse male fecundity and mating behavior, we used general linear mixed models (Bates and Maechler, 2009) and *lmertest* (Kuznetsova et al., 2017) to calculate *p*-values, tested with line fitted as a random effect. Degrees of freedom were based on the Satterthwaite approximation. In some cases, the response variable was square root transformed to satisfy model assumptions. For binomial data, *p*-values were calculated in *afex* (Singmann et al., 2016). All analyses were performed using R (v 3.4.0, R Core Team, 2017).

## RESULTS

### Sperm and SR length

Long sperm lines had significantly longer sperm than short sperm lines (long mean ± SE: 1934.00 ± 10.78, *n* = 86 sperm cells; short: 1673.97 ± 10.46, *n* = 115 sperm cells; *t*_192.94_ = 17.313, *P* = 2.2e-16). Likewise, long SR lines had significantly longer SRs than short SR lines (long: 2504.72 ± 58.74 µm, *n* = 22 SRs; short: 1640.56 ± 29.83 µm, *n* = 31 SRs; *t*_31.85_ = 13.17, *P* = 1.98e-14). However, sperm lengths were not significantly different between SR selection regimes (long: 1840.02 ± 6.83 µm, *n* = 119 sperm measurements; short: 1855.07 ± 6.76 µm, *n* = 117 sperm cells; *t_234_* = −1.567, *P* = 0.1185). Similarly, SR length in sperm selection treatments did not differ (long: 2138.50 ± 94.97µm, *n* = 6; short: 2237.69 ± 51.78 µm, *n* = 10; *t_8.02_* = −0.917, *P* = 0.39).

### Fitness

In the sperm selection lines, long sperm males had lower mating success (*χ*^2^ = 4.35, df = 1, *P* = 0.037; Fig 1a), suggesting that there is a pre-copulatory cost to the post-copulatory long sperm advantage. However, there were no differences in male attractiveness (mating latency; *F*_1,211_ = 2.270, *P* = 0.133; Fig 1c) or copulation duration (*F*_1,192_ = 0.553, *P* = 0.458; Fig 1e). Both males and females from long sperm lines trended toward higher fecundity, though this pattern was not statistically significant (males: *F*_1,5.8_ = 3.997, *P* = 0.094; Fig 2a; females: *F*_1,6_ = 3.560, *P* = 0.108; Fig 2c). We standardized fecundity within sex and selected trait (sperm or SR) by subtracting the mean and dividing the difference by the standard deviation, to directly compare fitness for both males and females (see Figure 3). Standardized fitness did not differ between males and females for short sperm (*F*_1,54.9_ = 0.119, *P* = 0.731) or long sperm lines (*F*_1,53.6_ < 0.001, *P* = 0.988; Fig. 3a). We did find a longevity advantage to long sperm genotypes in both sexes (males: *χ*^2^ = 32.50, df = 1, *p* = 0.001; sperm selected, females: *χ*^2^ = 9.13, df = 1, *P* = 0.003; Fig 4a, c). Higher survival specifically occurred for older females (Fig 4c) and at all ages for males (Fig 4a).

**FIGURE 1.**
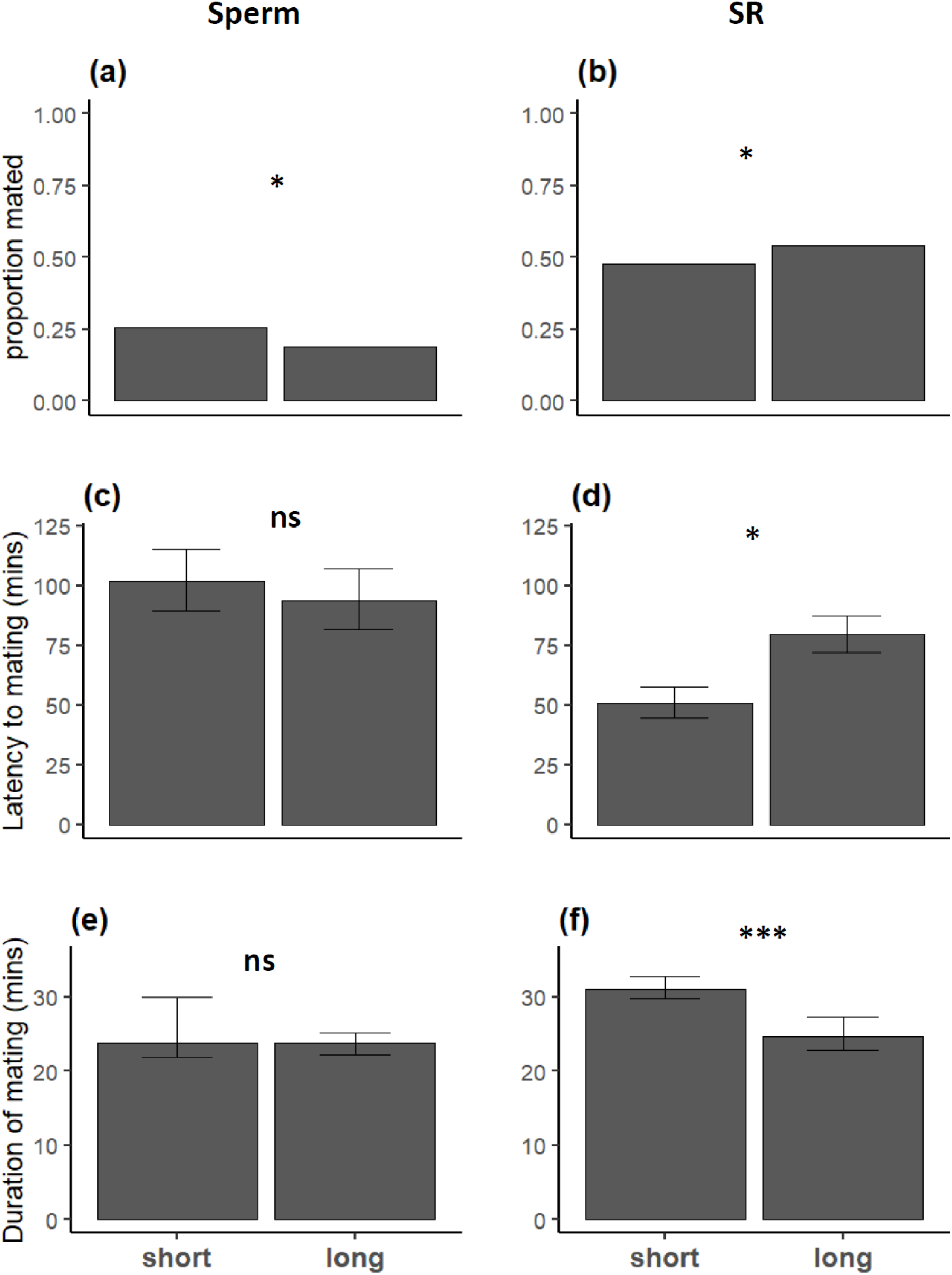
Mating success in males from the sperm selected (A, C, E) and SR selected (B, D, F) lines. Panels A) and B) show overall mating success (percentage of males that mated successfully). Mating latency, i.e. how long it took the successful males to start mating on average is shown in C) and D). Mating duration, shown in E) and F) indicates the mean duration of copulation within a treatment group.

**FIGURE 2:**
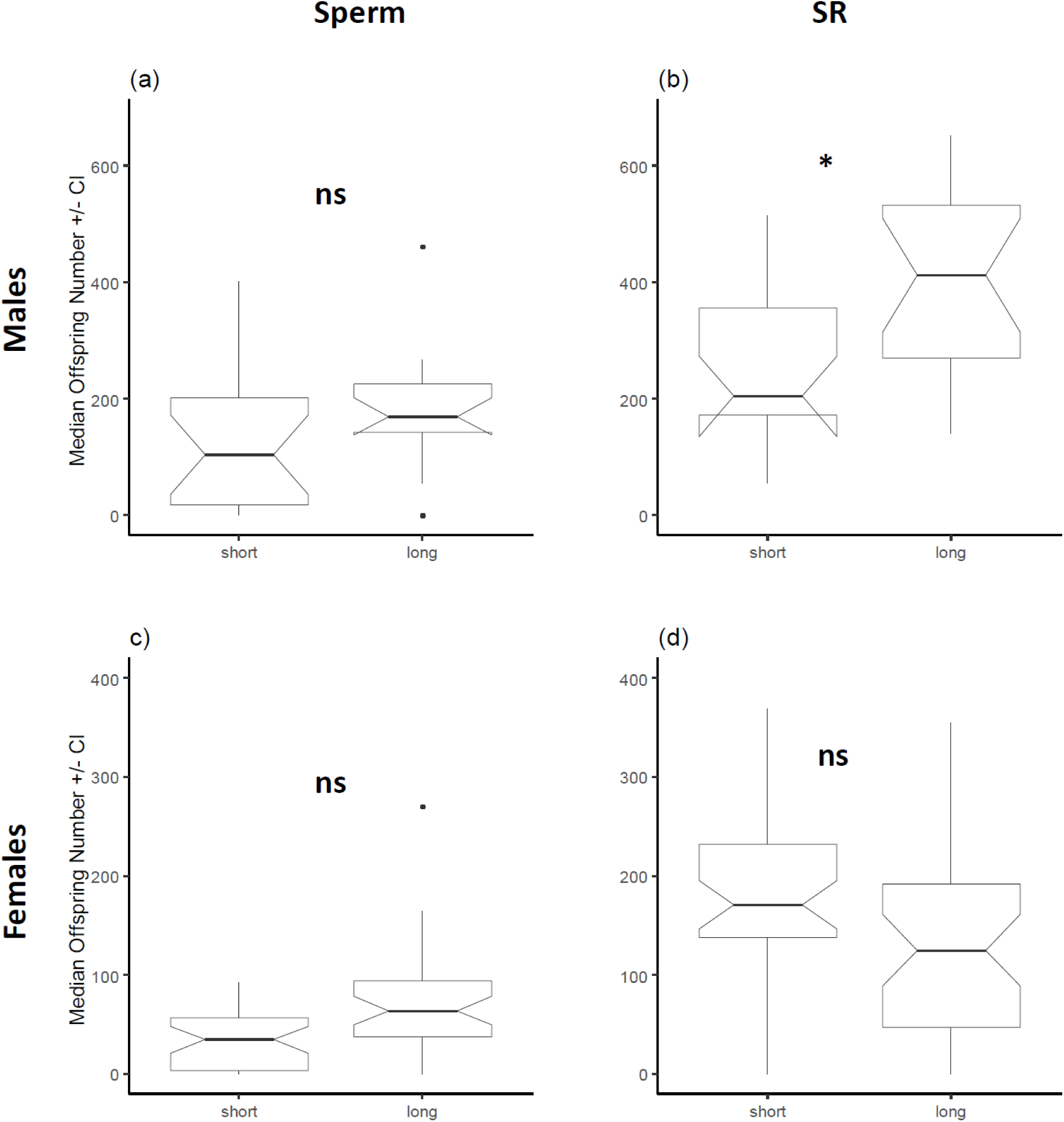
Number of offspring produced after mating trials by subsamples of males from long sperm (A), long SR (B), short sperm (C), and short SR (D) lines.

**FIGURE 3:**
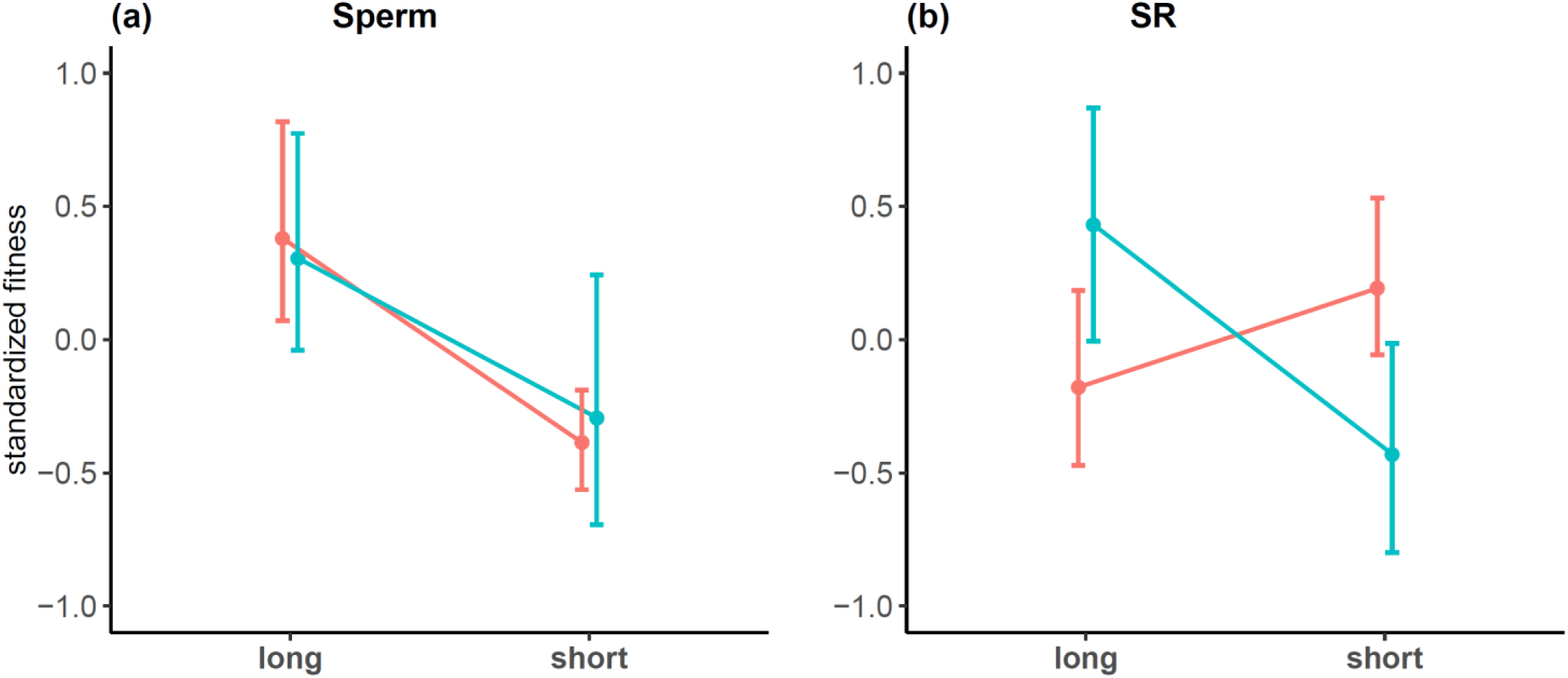
Standardized male and female fitness (mean ± bootstrapped 95% CI), for A) sperm and B) SR lines. Blue: short lines; red: long lines.

**FIGURE 4.**
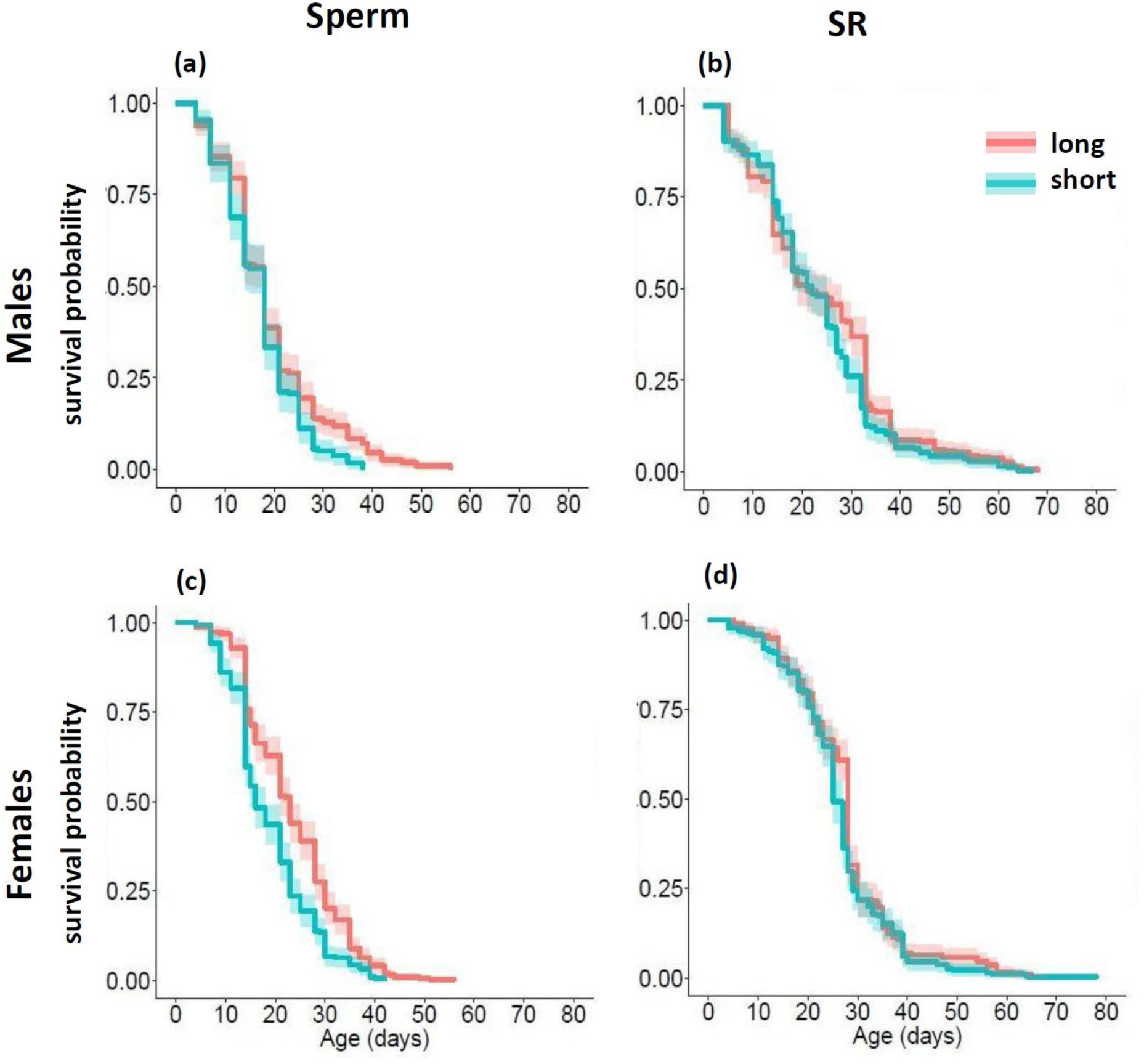
Survival curves of males from sperm selected lines (A) and of SR selected lines (B) and of females from sperm (C) and SR selected lines (D). Line colours represent selection regime (selection for short trait values: blue; long trait values: red). Shaded areas represent 95% confidence intervals. Age refers to adult age.

In the SR selection lines, short SR males were more attractive (shorter mating latency; *F*_1,569_ = 8.727, *P* = 0.003; Fig 1d) and copulated for longer (*F*_1,536_ = 91.261, *P* < 0.0001; Fig 1f), but long SR males ultimately had higher mating success (*χ*^2^ = 5.82, df = 1, *P* = 0.0158; Fig 1b) and higher fecundity (*F*_1,5.8_ = 6.118, *P* = 0.049, see Fig 2b). Females had higher relative fitness than males in short SR lines (*F*_1,52.4_ = 10.419, *P* = 0.002) and males had higher relative fitness than females in long SR lines (*F*_1,55.2_ = 7.485, *P* = 0.008; Fig. 3b). Interestingly, long SR females did not produce more offspring (*F*_1,6_ = 0.413, *P* = 0.544; Fig 2d), but they did live longer (*χ*^2^ = 4.64, df = 1, *p* = 0.031; Fig 4d), primarily at intermediate ages (Fig 4d). Male longevity was marginally longer between short and long selection regimes in SR selection lines (*χ*^2^ = 2.88, df = 1, *P* = 0.090; Fig 4b).

## DISCUSSION

Foundational questions in sexual selection ask how female preferences for elaborate male ornaments can evolve. That is, how do females benefit from these preferences, and what are the associated costs? There is ample evidence that, as predicted by theory (the Handicap Principle, Zahavi, 1975), ornaments are costly to produce and thus serve as signals of genetic quality (e.g., Godin and McDonough, 2003; Kotiaho, 2000; Manica et al., 2016; Mobley et al., 2018; Zuk et al., 1995). Females will gain indirect benefits from mating with high-condition males by having high-condition offspring (good genes; Fisher, 1958; Zahavi, 1977), if condition is heritable. If ornament phenotype is also heritable, females will additionally benefit by producing sexy sons, and if female preference is also heritable, a choosy female will have choosy daughters, who will also gain these indirect benefits.

On the other hand, intralocus conflict for either the trait that is exaggerated in males or its female preference (Lande, 1980; Rice, 1984) will constrain the evolutionary benefit of advantageous ornament or preference genotypes in males or females, respectively, by incurring fitness costs when those genotypes are expressed in the other sex (Chippindale et al., 2001; Cox and Calsbeek, 2009; Pischedda and Chippindale, 2006). Thus, the benefit of being a successful male may be limited by any costs of also having unfit daughters (Foerster et al., 2007). Likewise, any benefit of choosy daughters may be limited by low fitness of a female choice genotype in males.

In our study, genotypes producing long sperm or SRs confer multiple fitness benefits and few costs for both sexes (Table 1), suggesting that higher genetic quality is required to produce these traits. In particular, long selection lines for both sperm and SR phenotypes had increased longevity in males and females. By selecting for longer sperm, long SRs might also select for higher fitness genotypes in sons and daughters. Thus, the evolution of long sperm and long SRs may be driven by both viability selection (e.g., increased longevity) and indirect benefits (long SRs select for longer sperm, which confer fitness benefits to both sons and daughters). Together with a genetic correlation between the traits (Lüpold et al. 2006), these fitness benefits may aid in fueling a Fisherian runaway process. An alternate explanation for our results is that the selection and inbreeding history of the populations used in this experiment has led to the capture of genes conferring increased longevity in long sperm and long SR lines. It is important to note that increased longevity in both males and females is not necessarily indicative of increased lifetime reproductive success, which was not quantified here. Evaluation of fitness in unrelated populations with known sperm and SR phenotypes will be required to determine if sperm length and SR length are actually linked to “good genes”.

**Table 1.** Summary of results showing fitness benefits (+) or costs (−) of long phenotypes. Parentheses indicate a marginally insignificant trend, and NS indicates no significant difference between long and short phenotypes.

|  | Sperm |  | SR |  |
| --- | --- | --- | --- | --- |
|  | ♂ | ♀ | ♂ | ♀ |
| Mating success | - |  | + |  |
| Latency | NS |  | - |  |
| Copulation duration | NS |  | - |  |
| Fecundity | (+) | (+) | + | NS |
| Longevity | + | + | (+) | + |

We unexpectedly found that long SR genotypes in females confer increased longevity with no fecundity benefit, in contrast to previous work that showed that females with long SRs have higher reproductive output but at a cost to survival (Miller and Pitnick, 2003). These previous results may be due to increased storage capacity of both sperm and detrimental male ejaculate proteins (Chapman et al., 1995). In that study, long SRs were 40% longer than those reported here (3.5 mm vs 2.5 mm) and unlikely to occur naturally, perhaps because of these costs. Our more moderate SR lengths are comparable to those found in local wild *D. melanogaster* in the Washington, D.C. area (mean 2.5 mm, unpubl. data), and also on par with SR phenotypes shown to select for longer sperm (Miller and Pitnick, 2002). These moderately long SRs come with a longevity benefit, while also mediating sperm choice for longer sperm, perhaps reaping viability benefits for both sons and daughters. We thus find that both long sperm and long SRs may be honest signals of genetic condition.

Our results identified a tradeoff in males between long sperm and mating success, suggesting evolutionary modularity for traits under pre-copulatory versus post-copulatory sexual selection. In other words, long sperm confer only a post-copulatory advantage with no premating benefits. For males with long SR genotypes, reproductive success was mixed, with decreased attractiveness and copulation duration but increased mating success. This outcome may be due to more persistent courtship by long SR males, despite lower attractiveness, though we did not quantify courtship effort. At the same time, females mated to less attractive long SR males produced more progeny, suggesting a disconnect between male attractiveness and male fecundity. Higher fecundity in long SR males also further supports the hypothesis that genotypes associated with long SRs are of higher quality.

Most studies that examine the relationship between pre-copulatory and post-copulatory processes ask if mating success and attractiveness (pre-copulatory) is correlated with paternity outcome (post-copulatory). This study flips that question by starting with traits associated with paternity success (sperm and SR length) and looking for an association with premating outcome. We would not necessarily expect to find a difference between comparisons of pre-copulatory success with post-copulatory outcome, as opposed to associating post-copulatory outcome with pre-copulatory success. However, most studies in other species have found that pre-copulatory success is a good predictor of post-copulatory outcome (Evans et al., 2003; Hosken et al., 2008; Lewis and Austad, 1994; Polak and Simmons, 2009; Sbilordo & Martin, 2014; McDonald et al., 2017), though it matters which traits are considered (Ala-Honkola and Manier, 2016). Here, however, we did not find an association between sperm length and premating success, in concordance with Droge-Young et al. (2012) and Travers et al. (2016). It is possible that pre-copulatory and post-copulatory effort trade off in *D. melanogaster* (Filice and Dukas, 2019), such that males may invest in one or the other, but not both.

In conclusion, sperm length and SR length in this system do not appear to have fitness costs in the opposite sex. Rather, both long sperm and long SR phenotypes seem to confer fitness advantages to both males and females (with few costs), suggesting not only that long sperm are indeed an honest signal of good genes, but that female preference can also be an indicator of female quality. The costs and benefits incurred by female preferences have received less empirical attention than selection on male traits, primarily because female preferences (and concomitant costs and benefits) are more difficult to measure. Our work here suggests that selection driving male-female coevolution is not always antagonistic and can actually align to benefit both sexes.

## ACKNOWLEDGEMENTS

We are grateful to Zachary Boor, Michael West, and Ayesha Monga Kravetz for experimental assistance, and to Bob Cox for helpful discussion. (Financial disclosure statement removed for blinding purposes). We declare no conflicts of interest.

